# Plasma-driven biocatalysis using the cytochrome P450 enzyme CYP152_BSβ_

**DOI:** 10.1101/2025.01.09.631812

**Authors:** Tim Dirks, Sabrina Klopsch, Davina Stoesser, Sophie Desdemona Trenkle, Abdulkadir Yayci, Steffen Schüttler, Judith Golda, Julia Elisabeth Bandow

**Affiliations:** Applied Microbiology, Faculty of Biology and Biotechnology, Ruhr University Bochum, Germany; Plasma Interface Physics, Faculty of Physics and Astronomy, Ruhr University Bochum, Germany

**Keywords:** Cold atmospheric pressure plasma jet, cytochrome P450, immobilization, hydroxylation, decoy molecule

## Abstract

Plasma-driven biocatalysis utilizes *in situ* H_2_O_2_ production by atmospheric pressure plasmas to drive H_2_O_2_-dependent enzymatic reactions. Having previously established plasma-driven biocatalysis using recombinant unspecific peroxygenase from *Agrocybe aegerita* (r*Aae*UPO) to produce (*R*)-1-phenylethanol from ethylbenzene (ETBE), we here employed CYP152 from *Bacillus subtilis* (CYP152_BSβ_). CYP152_BSβ_ naturally hydroxylates medium and long-chain carboxylic acids, and, with short-chain carboxylic acids as decoy molecules, also converts non-natural substrates such as ETBE. To produce active CYP152_BSβ_ overexpression and heme loading were optimized. The conversion of the non-natural substrates guaiacol and ABTS with heptanoic acid as decoy molecule and H_2_O_2_ from stock solution yielded 18.28 and 21.13 nmol product min^-1^ 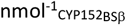, respectively. These reactions also served to assess compatibility of CYP152_BSβ_ with plasma-driven biocatalysis regarding temperature and H_2_O_2_ operating windows. To establish CYP152_BSβ_-based plasma-driven biocatalysis, immobilized enzyme in a rotating bed reactor (5 ml reaction volume) was then supplied with H_2_O_2_ from a capillary plasma jet operated with 1280 ppm H_2_O in helium. After a 120 min run time a turnover number (TON) of 18.82 mol_(*R*)-1-PhOl_ 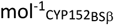 was reached. We conclude that plasma-driven biocatalysis can be extended to other H_2_O_2_-dependent enzymes. Future efforts will be directed at increasing the TON and product range.

## Introduction

Peroxidases and peroxygenases are promising enzyme classes for biocatalysis since they carry out one-electron oxidation reactions and stereoselective oxyfunctionalizations, which are difficult to perform by chemical means [1,2]. Cytochrome P450 (CYP450s) are peroxygenases in which the iron of the heme cofactor is typically ferric in the resting state with a characteristic absorbance at ∼418 nm (soret band) (reviewed in [3,4]). Upon binding carbon monoxide (CO), it forms a Fe^II^-CO complex and the maximum absorbance of the soret band shifts to 450 nm, hence the name of the enzyme class. P450 enzymes act as monooxygenases, catalyzing various reactions such as hydroxylations [5], epoxidations [6], reductive dehalogenation [7], sulfoxidation [8], and isomerizations [9]. P450 enzymes use *e*.*g*. NAD(P)H as an electron donor for their reaction cycle. In this cycle, the substrate binds to the active site of the enzyme, which results in the displacement of a water molecule. Upon the transfer of an electron originating from NAD(P)H to the active site of the enzyme, the formerly ferric heme iron is reduced to its ferrous state. Next, molecular oxygen binds covalently to the heme iron, and compound I is formed by a second electron transfer. The bound substrate is oxidized by two protons and one water molecule is released. Finally, oxidized substrate is released and H_2_O binds to the ferric heme iron, restoring the resting state of the enzyme [10].

The cytochrome P450 CYP152_BSβ_ of *Bacillus subtilis* belongs to a special subfamily that catalyzes the hydroxylation of long-chain carboxylic acids at the α- and β-position using H_2_O_2_ instead of NAD(P)H as electron donor [11,12]. This enzyme uses an alternative reaction pathway called peroxide shunt that by-passes some steps of the P450 enzyme reaction cycle [10,13]. Besides its natural activity, this enzyme oxidizes non-natural substrates when short chain carboxylic acids act as decoy molecules occupying the active site of the enzyme where they form a carboxylate-Arg^242^ salt bridge - a prerequisite enzyme activity [14,15]. The short-chain carboxylic acids are not long enough to be converted or to reach into the hydrophobic substrate channel. Instead, non-natural substrates can access the substrate channel where they get converted. It has previously been reported that the substrate ethylbenzene (ETBE) can be converted in an enantioselective reaction using the CYP152_BSβ_ enzyme, with an *ee* value of 68% for (*R*)-1-PhOl when heptanoic acid was used as a decoy molecule [10,14,15].

The aim of the present study was to establish CYP152_BSβ_-based plasma-driven biocatalysis. To this end, CYP152_BSβ_ was purified and enzyme kinetic parameters were investigated using the non-natural substrates guaiacol and ABTS and H_2_O_2_ supply from stock solutions. Using horseradish peroxidase (HRP), H_2_O_2_ it had been shown that H_2_O_2_ can be produced *in situ* using technical plasma to drive the reaction of H_2_O_2_-dependent enzymes [16] and an online biocatalysis has been established for recombinant unspecific peroxygenase from *Agrocybe aegerita* (r*Aae*UPO) [17].

### Experimental procedures

#### Plasma source and operating parameters

As plasma sources, the PlasmaDerm DBD (Cinogy, Germany) [18] and the atmospheric pressure capillary plasma jet [19,20] were used. The DBD had a copper electrode with a diameter of 20 mm, driven at V_RMS_ = 13.5 kV and a trigger frequency of 300 Hz. For plasma treatments, 40 µl samples were placed on grounded stainless-steel supports at a distance of 1 mm from the DBD.

The RF-driven atmospheric pressure capillary plasma jet with a plasma volume of 4 mm x 0.88 mm x 40 mm (outer capillary dimension 5 mm x 1.32 m x 40 mm) was used with an input power of 6.6 +/-0.6 W for plasma ignition. The gas flow (2 slm He) was split and (partially) routed through a bubbler with cooled deionized water to enrich the feed gas with water molecules. The distance between the nozzle of the capillary jet and the sample was approx. 16 mm. H_2_O_2_ production by this capillary plasma jet was previously characterized [20].

#### Plasmid construction

The *cypC* overexpression plasmid was designed based on pET28b using *Nde*I and *Hind*III restriction sites at the beginning and the end of the gene, respectively. The *B. subtilis* gene was codon optimized for *Escherichia coli* (*E. coli*). The vector-encoded C-terminal His-tag was silenced by preserving the natural stop codon of the *cypC* gene. Gene synthesis, cloning, and sequencing were performed by Genscript (USA). The plasmid enabling isopropyl-β-D-thiogalactopyranoside (IPTG)-dependent expression of the His_6_-*cypC* construct was transformed into chemically competent *E. coli* DH5α for amplification and then re-isolated. LB agar containing 50 µg ml^-1^ kanamycin was used for selection.

#### Overexpression and purification of CYP152_BSβ_

The plasmid pET28b::*cypC* was freshly transformed into chemically competent BL21 (DE3) *E. coli* cells. An overnight culture was prepared using LB medium supplemented with 50 µg ml^-1^ kanamycin for selection. For overproduction of CYP152_BSβ_ in LB medium, cultures (total volume of 1 L) were inoculated to an OD_600_ of 0.05 containing 50 µg ml^-1^ kanamycin, 200 µmol l^-1^ hemin chloride (stock solution dissolved in 100 mmol l^-1^ NaOH), and 500 µmol l^-1^ δ-aminolevulinic acid and incubated at 37°C to an OD_600_ of 0.5-0.6 followed by the addition of IPTG (100 µmol l^-1^). After 4 h incubation at 30°C, the cells were harvested for protein purification.

To improve heme loading, ZYM5052 auto-induction medium was used [21,22]. For the main culture, 1 L ZYM5052 was supplemented with 50 µg ml^-1^ kanamycin, 200 µmol l^-1^ hemin chloride (stock solution dissolved in 100 mmol l^-1^ NaOH), and 500 µmol l^-1^ δ-aminolevulinic acid and inoculated with 10 ml of a preculture. Overexpression was allowed to proceed for 5 days at 16°C and 120 rpm. Every 24 h, samples were withdrawn for SDS-PAGE and western blot analysis according to standard protocols and cell densities of the samples were normalized. His_6_-tagged proteins were detected using a fluorescence-based 6 x His-tag monoclonal antibody (ThermoFisher Scientific, USA) and a ChemiDoc^TM^ MP Imaging System (Bio-Rad, USA). After 5 days, cells of the main culture were harvested, washed with 100 mmol l^-1^ potassium phosphate buffer (pH 7), and stored at -80°C until further use.

For cell disruption, cells were resuspended in lysis buffer (0.2 mg ml^-1^ DNase, 0.2 mg ml^-1^ RNase, 0.35 mg ml^-1^ lysozyme, 2 mmol l^-1^ DTT, and cOmplete protease inhibitor (Roche, Switzerland) in 100 mmol l^-1^ potassium phosphate buffer containing 300 mmol l^-1^ potassium chloride and 20% glycerol, pH 7). After cell disruption using a pressure cell homogenizer (FPG12800, Homogenising Systems, United Kingdom) for six cycles and centrifugation for 30 min (21.000 g), the supernatant was loaded onto a HisTrap FF crude 5 ml column (GE Healthcare, USA) and the His_6_-tagged CYP152_BSβ_ protein was purified with an ÄKTA pure25 system (GE Healthcare, USA). Proteins were eluted with three stepwise increases in imidazole concentration using three column volumes (50 mmol l^-1^; 75 mmol l^-1^, 200 mmol l^-1^). The HisTrap FF crude 5 ml column was finally washed with 500 mmol l^-1^ imidazole for ten column volumes to remove remaining bound protein. Elution fractions of interest were pooled, concentrated using centrifugal filter units (10 kDa molecular weight cut-off), and used for reconstitution.

#### Reconstitution

To ensure sufficient heme loading, the protein concentration of purified CYP152_BSβ_ was determined using the Bradford method and incubated with a two-fold molar ratio of hemin chloride (in a total volume of approx. 50 ml) at 25°C for 1 h. Unbound hemin chloride was removed by overnight dialysis against 100 mmol l^−1^ potassium phosphate buffer containing 300 mmol l^-1^ potassium chloride and 20% glycerol, pH 7. The dialyzed and reconstituted CYP152_BSβ_ was aliquoted and stored at -80°C.

#### Spectral analysis

Absorption spectra of CYP152_BSβ_ were recorded using an enzyme concentration of 12 µmol l^-1^ in a total volume of 100 µl (100 mmol l^−1^ potassium phosphate buffer containing 300 mmol l^-1^ potassium chloride and 20% glycerol, pH 7) in a UV/VIS spectrophotometer (V-750 spectrophotometer, Jasco, Germany). Buffer was used as blank. R/z values were calculated by relating absorption of the Soret peak at its maximum intensity (∼420 nm) to absorption at 280 nm [23,24].

#### Activity assays

Enzyme activity of CYP152_BSβ_ was determined based on the conversion of the natural substrate myristic acid to α- and β-hydroxylated myristic acid. To this end, 120 µmol l^-1^ myristic acid was mixed with 0.1 µmol l^-1^ CYP152_BSβ_ in potassium phosphate buffer (100 mmol l^-1^, pH 7) in a total volume of 200 µl. To start the reaction, 0.5 mmol l^-1^ H_2_O_2_ was added and the reaction was allowed to proceed at 37°C for 30 min under constant shaking (500 rpm). Afterwards, analysis was performed according to Girhard *et al*. [25]. Briefly, samples were mixed twice with 500 µl diethyl ether and centrifuged. The organic phase was transferred into a new vessel and dried using anhydrous MgSO_4_. The supernatant was again transferred into a new vessel followed by evaporation of the organic phase at 34.6°C for 20 min. Residual was resuspended using 50 µl N,O-bis(trimethylsilyl)trifluoroacetamide with 1% trimethylchlorosilane (BSTFA-TMCS, TCI Chemicals, Germany) followed by an incubation at 80°C for 30 min. Derivatized samples were immediately measured by gas chromatography (Shimadzu GC-2030 Nexis, Japan) along with EI-MS using a GC/MS-QP2020 NX (Shimadzu, Japan) equipped with a FS-Supreme-5ms column (30 m x 0.25 mm x 0.25 µm, Chromatographie Service, Germany).

Column temperature was held at 160°C for 1 min and ramped to 260°C with a velocity of 10°C min^-1^. Finally, temperature was increased to 300°C (40°C min^-1^) and held for 3 min. Helium was used as the carrier gas with a flow rate of 1 ml min^-1^. For mass spectrometric analysis, mass detector was operated in electron impact (EI) mode at 70 eV with an electron multiplier voltage of 1.25 kV. Sizes of obtained mass fragments were compared to Girhard *et al*. [25].

Conversion of non-natural substrates was analyzed using 2,2’-azino-bis(3-ethylbenzothiazoline-6-sulfonic acid) (ABTS) or guaiacol together with heptanoic acid as a decoy molecule according to Shoji and Watanabe [10]. Activity assays were performed following a basic protocol with slight modifications for decoy molecule addition. 5 mmol l^−1^ 2,2′-azino-bis(3-ethylthiazoline-6-sulfonate) (ABTS; ε = 36.8 mmol l^−1^ cm^−1^) or 50 mmol l^−1^ guaiacol (ε = 26.6 mmol l^−1^ cm^−1^) were used. The substrates were dissolved in 100 mmol l^−1^ sodium citrate buffer (pH 5) and 100 mmol l^-1^ potassium phosphate buffer (containing 300 mmol l^-1^ potassium chloride and 20% glycerol, pH 7), respectively, while 20 mmol l^-1^ heptanoic acid (dissolved in EtOH) was added. Reactions were initiated by always adding the same volume of H_2_O_2_ solution prepared by dilution in distilled water (*A. dest*.) to yield concentrations ranging from 0 to 10 mmol l^-1^ and product formation was monitored at 405 nm or 470 nm for 2 min using a microplate reader (Biotek, Epoch, Germany). The enzyme concentration was 1 μmol l^-1^. Activity was calculated from the initial reaction velocity within the first seconds of the kinetic (formula: Δabsorption/Δtime).

The temperature optimum was tested in a range of 10 to 40°C in 10°C increments using a UV/VIS spectrophotometer with built-in Peltier element (V-750 spectrophotometer, Jasco) in a total volume of 100 µl. All reaction components were incubated at the respective temperature for 10 min before starting the reaction.

#### Plasma-driven biocatalysis

Immobilization with ReliZyme HA403 M beads was carried out as described previously [17] by applying 10 µmol l^-1^ CYP152_BSβ_ to 200 mg HA403 M beads in a volume of 5 ml 100 mmol l^−1^ potassium phosphate buffer containing 300 mmol l^-1^ potassium chloride and 20% glycerol (pH 7). Binding efficiency was determined by comparing enzyme activity of the supernatant to enzyme activity of the solution initially used for immobilization. Plasma-driven biocatalysis was performed as described previously for the recombinant unspecific peroxygenase of *Agrocybe aegerita* r*Aae*UPO [17]. In short, protein-loaded beads (100 mg) were transferred to a rotating bed reactor (build in-house, dimensions: Ø2 cm x 0.7 cm). The reactor was placed in a vessel filled with 5 ml potassium phosphate buffer (100 mmol l^-1^, pH 7) containing 50 mmol l^-1^ ethylbenzene (ETBE) and 20 mmol l^-1^ heptanoic acid (dissolved in EtOH). Plasma treatment was performed for up to 60 min, and 150 µl-samples were withdrawn every 5 min for analysis of product formation [16,17]. To compensate the sample withdrawal and evaporation, 300 µl of potassium phosphate buffer (100 mmol l^-1^, pH 7) containing 50 mmol l^-1^ ETBE and 20 mmol l^-1^ heptanoic acid were resupplied.

To avoid H_2_O_2_ accumulation, the entire reaction solution was exchanged every 10 min. Product accumulation in the reaction volume was measured by GC [16,17]. Plasma treatment was continued for a total of 120 min.

## Results and Discussion

In an initial attempt to produce CYP152_BSβ_ for plasma-driven biocatalysis, we overexpressed CYP152_BSβ_ using LB medium as described in the methods section. While we obtained satisfactory amounts of protein, heme loading of the enzyme was poor reaching R/z values (ratio A_420_/A_280_ nm) of 0.3 (Supplementary Figure 1). To improve heme loading and protein folding we used a recently published overexpression protocol for recombinantly expressed peroxygenases, which utilized an auto-induction medium [22]. We overexpressed *cypC* for five days at 16°C and purified CYP152_BSβ_ using an IMAC column (Supplementary Figure 2). SDS-PAGE and western blot analysis revealed that the fraction containing the first elution peak contained impurities, while the purity of the pooled fraction containing peaks two and three (CYP152_BSβ_) was very high (Figure 1). Additionally, CYP152_BSβ_ was detected in the pellet after cell lysis hinting at formation of inclusion bodies, and in the flowthrough, hinting at an overloading of the IMAC column. Despite this loss, we obtained large quantities of pure protein (350 mg_protein_ l^-1^_culture_ yield). To maximize heme loading, hemin chloride was added to the purified protein to allow heme-free CYP152_BSβ_ to incorporate heme. This reconstitution step resulted in an increase in R/z value from 1.01 before to 1.85 after reconstitution (Figure 2).

**Figure 1:**
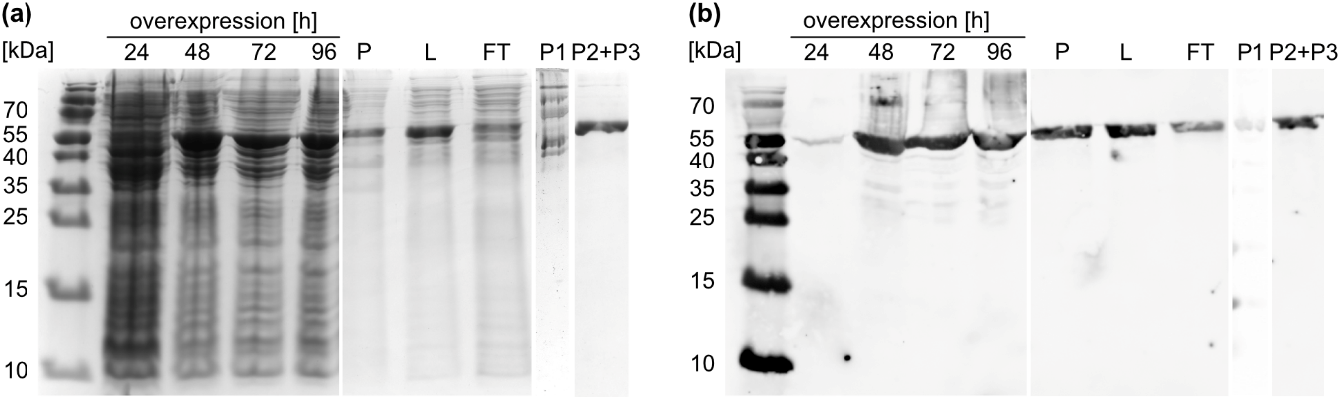
SDS-PAGE (a) and western blot (b) analysis of overexpression and IMAC purification of CYP152_BSβ_. The enzyme was purified from lysate as described in the method section. Samples of overexpression after 24 – 96 h (lanes 2-5; density was adjusted by adding 75 µl SDS loading buffer to 1 ml cell pellet with an OD_600_ of 0.3), pellet after lysis (P, 1:20 dilution), lysate (L, 1:10 dilution), flowthrough (FT, 1:10 dilution), fraction containing the first peak of the IMAC purification (P1, 5 µg), and purified protein (P2+P3, pooled fraction containing peaks two and three, 5 µg) were subjected to denaturing SDS-PAGE using 10 µl protein solution per lane. The expected molecular weight of CYP152_BSβ_ was approx. 50.5 kDa.

**Figure 2:**
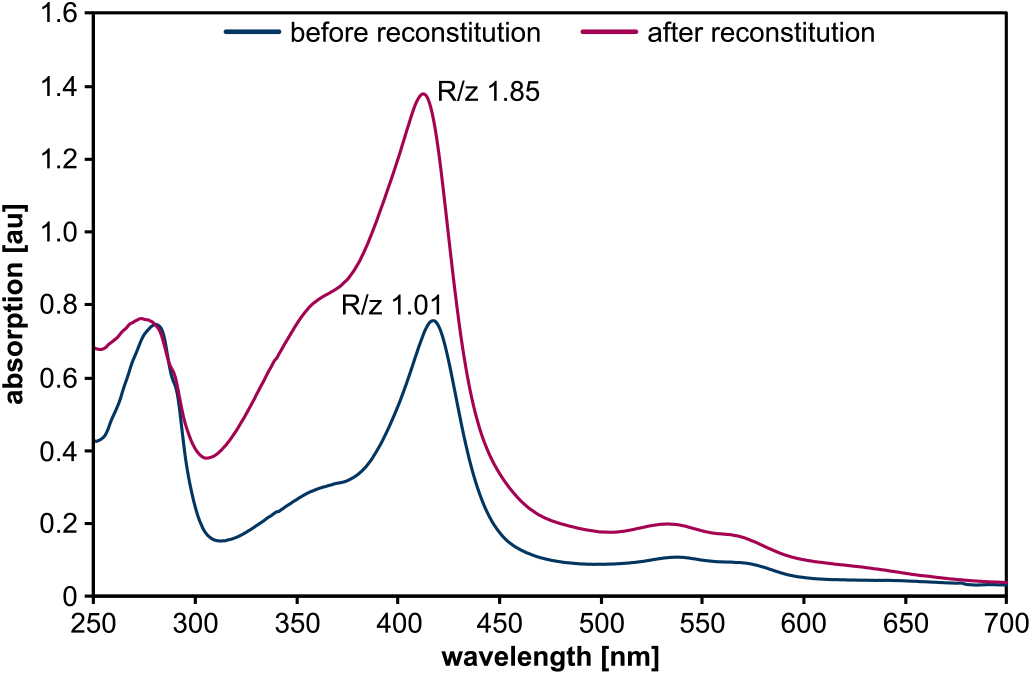
Spectral analysis of purified CYP152_BSβ_ before and after reconstitution. Absorption spectra were recorded using 12 µmol l^-1^ CYP152_BSβ_ in potassium phosphate buffer (100 mmol l^-1^, pH 7). Buffer served as blank. R/z values were calculated by relating absorption of the soret peak (at 420 nm) to absorption at 280 nm. Data shown are representative of three independent replicates.

The purified CYP152_BSβ_ enzyme was tested for catalytic activity using the conversion of myristic acid to (α)- and (β)-hydroxy myristic acid supplying the co-substrate H_2_O_2_ from a stock solution (Supplementary Figure 3) [11]. The enzyme converted myristic acid to the products α- and β-hydroxy myristic acid, which were identified by GC/MS analysis based on their parental masses and described fragmentation patterns (Supplementary Figure 4) [11]. The observed product distribution of 40% (α)-hydroxy myristic acid : 60% (β)-hydroxy myristic acid is well in line with previous reports [13].

Beside its natural substrates, CYP152_BSβ_ can convert non-natural substrates when short-chain carboxylic acids serve as decoy molecules to engage the active site. Among the non-natural substrates converted is ethylbenzene. Its conversion to 1-phenylethanol is the model reaction of the unspecific peroxygenase from *Agrocybe aegerita* (*Aae*UPO) which was previously used in plasma-driven biocatalysis [16,17]. In previous reports, heptanoic acid served as a decoy molecule resulting in comparatively high activities and good stereoselectivity (*ee* (*R*)-1-PhOl of approx. 70%) [15]. To record Michaelis-Menten kinetics we focused on heptanoic acid as decoy molecule, combining it with the conversion of the non-natural substrates guaiacol and ABTS whose conversion can easily be monitored photometrically (Figure 3, Table 1). Using H_2_O_2_ from stock solutions, K_M_ values of 0.25 (ABTS) and 0.57 mmol l^-1^ H_2_O_2_ (guaiacol) and corresponding turnover numbers of 21.13 to 18.28 nmol product min^-1^ 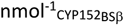 were obtained, respectively.

**Figure 3:**
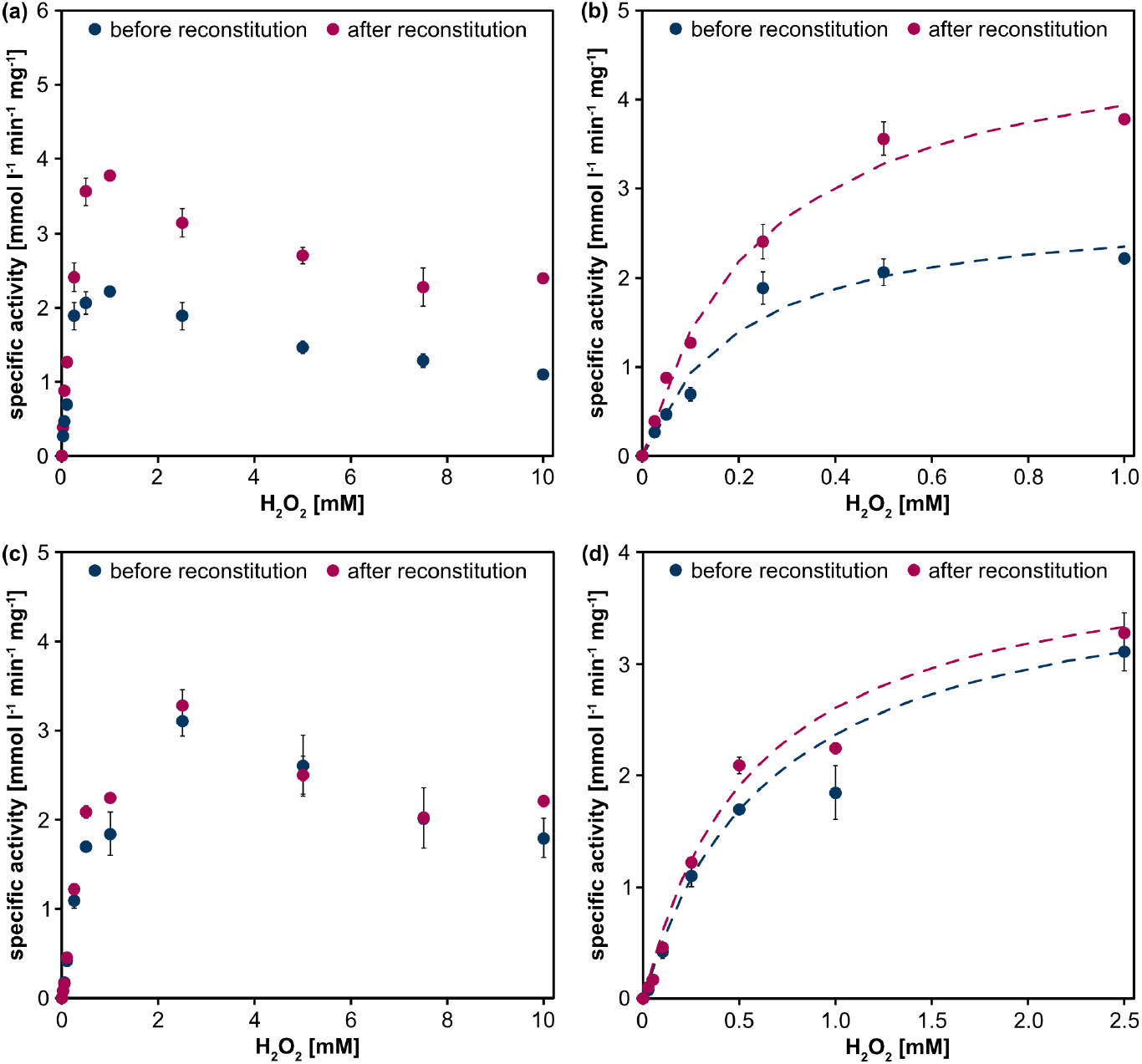
Activity of CYP152_BSβ_ at different H_2_O_2_ concentrations using the non-natural substrates ABTS (a,b) and guaiacol (c,d). Activity assays were performed at room temperature (approx. 22°C) using heptanoic acid as decoy molecule. Product conversion was measured photometrically at 405 nm (ABTS) or 470 nm (guaiacol) and product formation was calculated using Lambert-Beer law (ε_ABTS_ 36.8 mmol l^-1^ cm^-1^; ε_guaiacol_ 26.6 mmol l^-1^ cm^-1^). Specific activities were calculated based on the initial reaction velocities and given in mmol l^-1^ min^-1^ mg^-1^. Reactions were performed by adding up to 10 mM H_2_O_2_. For the conversion of ABTS, specific activity peaked at 1 mM, for guaiacol it peaked at 2.5 mM ((b,d) show blow-ups of the activity at low H_2_O_2_ concentrations, the dashed lines indicate the H_2_O_2_-dependent increase in specific activity). Means and standard deviations represent three independent experiments.

**Table 1:** Kinetic parameters of CYP152_BSβ_. Calculations were performed based on the data shown in Figure 3.

| | $K_M (H_2O_2) [mmol\ l^{-1}]$ | | $nmol\ product\ min^{-1}\ nmol^{-1}\ CYP152_{BS\beta}$ | |
| --- | --- | --- | --- | --- |
|  | before reconstitution | after reconstitution | before reconstitution | after reconstitution |
| ABTS | $0.20 \pm 0.16$ | $0.25 \pm 0.11$ | $12.41 \pm 0.12$ | $21.13 \pm 0.31$ |
| Guaiacol | $0.66 \pm 0.21$ | $0.57 \pm 0.23$ | $17.69 \pm 0.99$ | $18.28 \pm 0.16$ |

The influence of the temperature on CYP152_BSβ_ activity was evaluated using ABTS and guaiacol as substrates (Figure 4). Increasing temperatures led to higher activities using both substrates with ABTS conversion peaking at 30°C (18.62 mmol l^-1^ min^-1^ mg^-1^) and guaiacol conversion increasing to 17.27 mmol l^-1^ min^-1^ mg^-1^ at 40°C. As was to be expected, higher specific activities were reached for the reconstituted compared to the crude CYP152_BSβ_. Generally, activity increased ∼7.5x when the temperature increased from 10°C to 40°C. At temperatures > 40°C the enzyme precipitated. Thus, working ranges from 11.04 to 18.62 mmol product min^-1^ 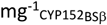 (ABTS) and 2.27 to 17.27 mmol product min^-1^ 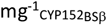 were identified.

**Figure 4:**
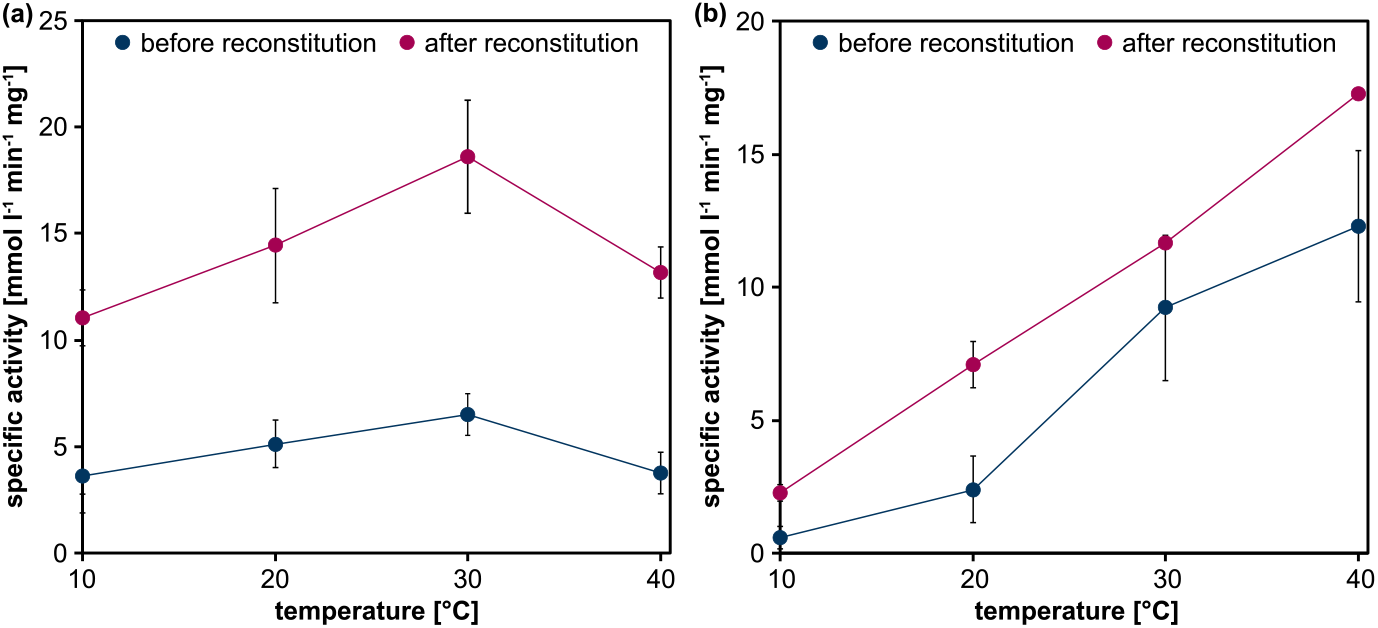
Temperature dependency of CYP152_BSβ_ using ABTS (a) and guaiacol (b) as non-natural substrates. Product conversion was measured photometrically at 405 nm or 470 nm, respectively. Activity assays were performed at different temperatures using a UV/VIS spectrometer with an integrated Peltier element. All assay components were incubated at the respective temperatures for 5 min prior to starting the enzymatic reaction. Product formation was calculated using Lambert-Beer law (ε_ABTS_ 36.8 mmol l^-1^ cm^-1^; ε_guaiacol_ 26.6 mmol l^-1^ cm^-1^). Specific activities are given in mmol l^-1^ min^-1^ mg^-1^ and were calculated based on the initial velocities of the reactions. Means and standard deviations reflect three experiments.

After confirming enzyme activity of CYP152_BSβ_, its application in plasma-driven biocatalysis was assessed. Non-thermal plasma is used to generate H_2_O_2_ *in situ* allowing to adjust the H_2_O_2_ production level specifically to the needs of H_2_O_2_-dependent enzymes [16,17]. In particular, accumulation of H_2_O_2_ can be avoided, preventing inactivation of these heme-containing enzymes [26,27]. To date, such an online plasma-driven biocatalysis has only been established for the recombinant unspecific peroxygenase from *Agrocybe aegerita* [17]. In a first attempt to realize plasma-driven biocatalysis with CYP152_BSβ_, we turned to the conversion of ETBE to (*R*)-1-PhOl, the reaction established for plasma-driven biocatalysis with r*Aae*UPO. To modulate H_2_O_2_ production by the capillary plasma jet, water concentrations of 1280 and 6400 ppm H_2_0 were used in the feed gas. 6400 ppm H_2_O water in the feed gas led to a rapid inactivation of the enzyme (Supplementary Figure 5), presumably due to the accumulation of excess H_2_O_2_. Based on previous analyses it is expected that within a 5 min treatment the capillary plasma jet supplies the reaction volume with >0.7 mM H_2_O_2_ for enzymatic conversion [28]. However, with a water concentration of 1280 ppm H_2_O in the feed gas supplying approximately 0.25 mM H_2_O_2_ in 5 min, in the tested reaction scheme plasma-driven biocatalysis with reconstituted CYP152_BSβ_ led to a constant accumulation of product reaching 60 µmol l^-1^ after 40 min reaction time (Figure 5).

**Figure 5:**
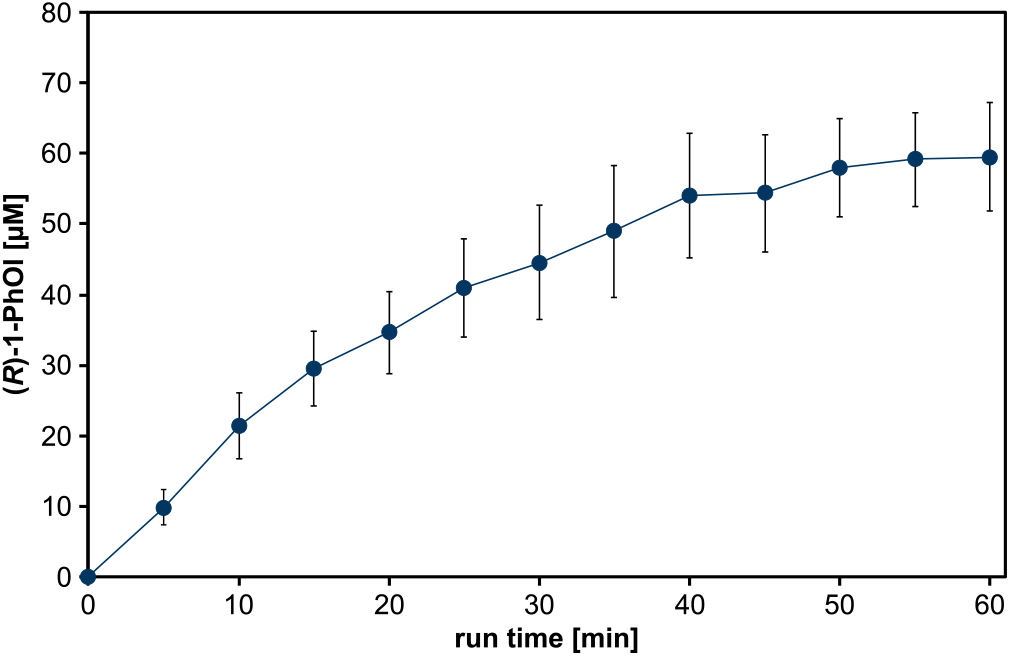
Plasma-driven biocatalysis (1280 ppm H_2_O in feed gas) with capillary plasma jet using CYP152_BSβ_. Product conversion of the substrate ETBE using direct plasma treatment of CYP152_BSβ_ immobilized on ReliZyme HA403 M beads. Reaction solution contained 5 ml potassium phosphate buffer (100 mmol l^-1^, pH 7) with 50 mmol l^-1^ ETBE and 20 mmol l^-1^ heptanoic acid (as decoy molecule). Plasma treatment was performed with a water concentration of 1280 ppm in the feed gas. Every 5 min, aliquots were withdrawn for product analysis by GC measurement and the reaction volume was topped up with reaction solution to 5 ml. Means and standard deviations reflect three experiments.

As expected for enzyme-based catalysis, the enantioselectivity of the reaction, previously described with an *ee* value of 68%, was preserved when H_2_O_2_ was generated using plasma (Supplementary Figure 6). After 40 min production ceased and at 60 min residual enzyme activity was shown to be drastically reduced (∼20%, Supplementary Table 1). Considering the low conversion rates, the enzyme potentially still suffered from excess H_2_O_2_-induced inactivation. In an attempt to increase enzyme lifetime, the H_2_O concentration in the feed gas was further lowered to 640 ppm. This, however, resulted in decreased product formation rates (Supplementary Figure 7). Reverting to 1280 ppm H_2_O in the feed gas, we then tested a different reaction scheme recently described [17], in which the complete reaction solution was exchanged every 10 min, thereby replenishing substrate and eliminating H_2_O_2_ and product (Figure 6).

**Figure 6:**
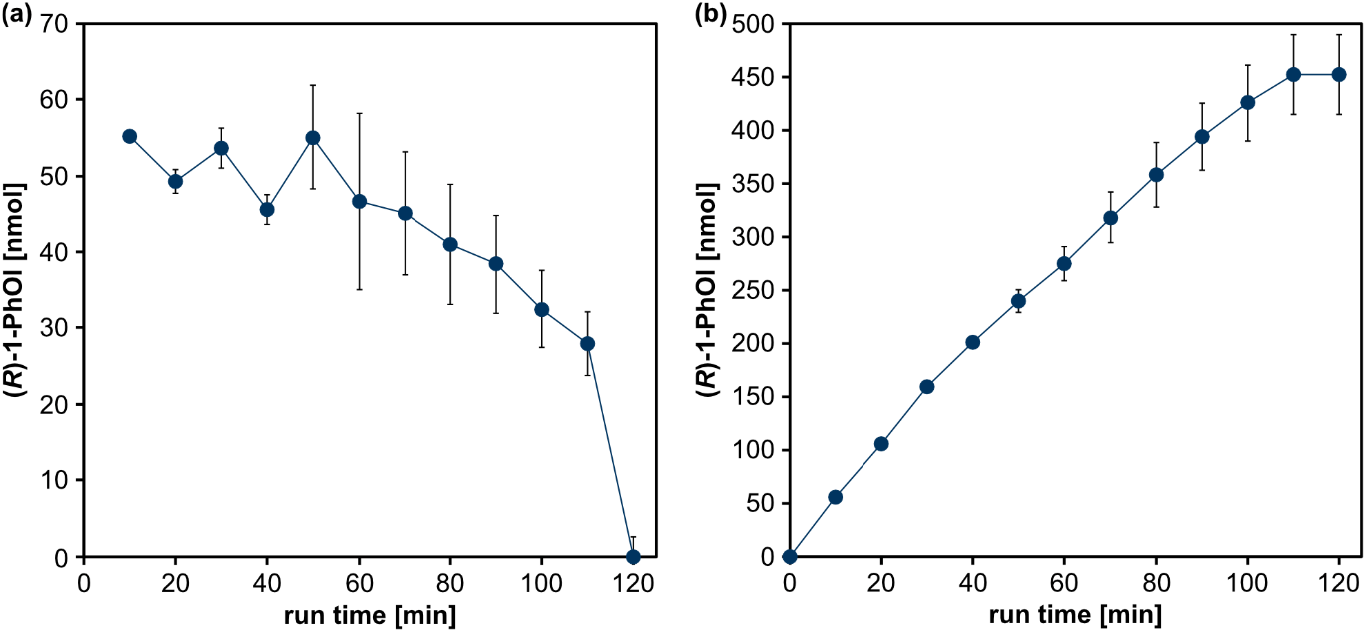
Plasma-driven biocatalysis with the capillary plasma jet using CYP152_BSβ_ using complete buffer exchange every 10 min. The reaction solution containing 5 ml potassium phosphate buffer (100 mmol l^-1^, pH 7) with 50 mmol l^-1^ ETBE and 20 mmol l^-1^ heptanoic acid (as decoy molecule) and was exchanged every 10 min. Product conversion of the substrate ETBE were evaluated every 10 min (a) and product accumulation (b) were evaluated with H_2_O_2_, which was supplied by plasma treatment of the reaction solution containing CYP152_BSβ_ immobilized on ReliZyme HA403 M using a capillary plasma jet and 1280 ppm H_2_O in the feed gas by GC measurement. Plasma treatment was continued for a total of 120 min. Means and standard deviations reflect three experiments.

Product formation using the frequent buffer exchange was stable during the first 50 min of the process, ranging from 45 to 55 nmol per cycle (Figure 6a). Enzyme inactivation set in at 60 min and reached approx. 100% after a run time of 120 min. Using this operating scheme, a total of 453 nmol of (*R*)-1-PhOl were produced and a turnover number (TON) of 18.82 mol_(*R*)-1-PhOl_ mol^-1^__CYP152BSβ__ was reached (Figure 6b). These values are well in line with the previously reported catalytic activity of 28 nmol_(*R*)-1-PhOl_ nmol^-1^__CYP152BSβ__ with H_2_O_2_ supplied from stock solutions [14]. Taken together, we showed that is suited for plasma-driven biocatalysis. r*Aae*UPO is the superior enzyme for the model ETBE hydroxylation reaction, but the use of enzymes like CYP152_BSβ_ could allow an expansion of the substrate and product spectrum of plasma-driven biocatalysis.

## Author’s contributions

T.D.: conceptualization, formal analysis, investigation, visualization, writing of original draft; S.K.: investigation, writing – review and editing; D.S.: investigation; S.D.T: investigation; A.Y.: conceptualization; S.S.: resources; J.G.: resources, writing – review and editing; J.E.B.: conceptualization, funding acquisition, supervision, writing – review and editing.

## References

[1] S. Bormann, A.G. Baraibar, Y. Ni, D. Holtmann, F. Hollmann, Specific oxyfunctionalisations catalysed by peroxygenases: opportunities, challenges and solutions, Catal Sci Technol. 5 (2015) 2038–52. doi: 10.1039/c4cy01477d.

[2] Y. Wang, D. Lan, R. Durrani, F. Hollmann, Peroxygenases en route to becoming dream catalysts. What are the opportunities and challenges? Curr Opin Chem Biol. 37 (2017) 1–9. doi: 10.1016/j.cbpa.2016.10.007.

[3] S.L. Kelly, D.E. Kelly, Microbial cytochromes P450: biodiversity and biotechnology. Where do cytochromes P450 come from, what do they do and what can they do for us? Philos Trans R Soc Lond B Biological Sci. 368 (2013) 20120476. doi: 10.1098/rstb.2012.0476.

[4] L. Hammerer, Regioselective biocatalytic hydroxylation of fatty acids by cytochrome P450s, Catal Lett. 148 (2018) 787–812. doi: 10.1007/s10562-017-2273-4.

[5] C.J.C. Whitehouse, S.G. Bell, L.-L. Wong, P450 BM3 (CYP102A1): Connecting the dots, Chem Soc Rev. 41 (2011) 1218–60. doi: 10.1039/c1cs15192d.

[6] E.H. Oliw, J. Bylund, C. Herman, Bisallylic hydroxylation and epoxidation of polyunsaturated fatty acids by cytochrome P450, Lipids. 31 (1996) 1003–21. doi: 10.1007/bf02522457.

[7] H.J. Ahr, L.J. King, W. Nastainczyk, V. Ullrich, The mechanism of reductive dehalogenation of halothane by liver cytochrome P450, Biochem Pharmacol. 31 (1982) 383–90. doi: 10.1016/0006-2952(82)90186-1.

[8] T.J. Volz, D.A. Rock, J.P. Jones, Evidence for two different active oxygen species in cytochrome P450 BM3 mediated sulfoxidation and N-dealkylation reactions, J Am Chem Soc. 124 (2002) 9724–25. doi: 10.1021/ja026699l.

[9] L. Gao, Y. Tu, P. Wegman, S. Wingren, L.A. Eriksson, A mechanistic hypothesis for the cytochrome P450-catalyzed cis–trans isomerization of 4-hydroxytamoxifen: An unusual redox reaction, J Chem Inf Model. 51 (2011) 2293–2301. doi: 10.1021/ci2001082.

[10] O. Shoji, Y. Watanabe, Monooxygenation of nonnative substrates catalyzed by bacterial cytochrome P450s facilitated by decoy molecules, Chem Lett. 46 (2017) 278–88. doi: 10.1246/cl.160963.

[11] I. Matsunaga, A. Ueda, N. Fujiwara, T. Sumimoto, K. Ichihara, Characterization of the ybdT gene product of Bacillus subtilis: Novel fatty acid β-hydroxylating cytochrome P450, Lipids. 34 (1999) 841–46. doi: 10.1007/s11745-999-0431-3.

[12] F.P. Guengerich, A.W. Munro, Unusual cytochrome P450 enzymes and reactions, J Biol Chem. 288 (2013) 17065–73. doi: 10.1074/jbc.r113.462275.

[13] T. Fujishiro, O. Shoji, S. Nagano, H. Sugimoto, Y. Shiro, Y. Watanabe, Crystal structure of H_2_O_2_-dependent cytochrome P450_SPα_ with its bound fatty acid substrate, J Biol Chem. 286 (2011) 29941–50. doi: 10.1074/jbc.m111.245225.

[14] O. Shoji, T. Fujishiro, H. Nakajima, M. Kim, S. Nagano, Y. Shiro, Y. Watanabe, Hydrogen peroxide dependent monooxygenations by tricking the substrate recognition of cytochrome P450_BSβ_, Angew Chem Int Ed. 46 (2007) 3656–59. doi: 10.1002/anie.200700068.

[15] O. Shoji, Y. Watanabe, Design of H_2_O_2_-dependent oxidation catalyzed by hemoproteins, Metallomics. 3 (2011) 379–88. doi: 10.1039/c0mt00090f.

[16] A. Yayci, Á.G. Baraibar, M. Krewing, E.F. Fueyo, F. Hollmann, M. Alcalde, R. Kourist, J.E. Bandow, Plasma-driven in situ production of hydrogen peroxide for biocatalysis, ChemSusChem. 13 (2020) 2072–79. doi: 10.1002/cssc.201903438.

[17] A. Yayci, T. Dirks, F. Kogelheide, M. Alcalde, F. Hollmann, P. Awakowicz, J.E. Bandow, Microscale atmospheric pressure plasma jet as a source for plasma-driven biocatalysis, ChemCatChem. 12 (2020) 5893–97. doi: 10.1002/cctc.202001225.

[18] S. Baldus, D. Schröder, N. Bibinov, V.S. der Gathen, P. Awakowicz, Atomic oxygen dynamics in an air dielectric barrier discharge: a combined diagnostic and modeling approach, J Phys D Appl Phys. 48 (2015) 275203–14. doi: 10.1088/0022-3727/48/27/275203.

[19] T. Winzer, D. Steuer, S. Schüttler, N. Blosczyk, J. Benedikt, J. Golda, RF-driven atmospheric-pressure capillary plasma jet in a He/O_2_ gas mixture: Multi-diagnostic approach to energy transport, J Appl Phys. 132 (2022) 183301. doi: 10.1063/5.0110252.

[20] S. Schüttler, A.L. Schöne, E. Jeß, A.R. Gibson, J. Golda, Production and transport of plasma-generated hydrogen peroxide from gas to liquid, Phys Chem Chem Phys. 26 (2024) 8255–8272. doi: 10.1039/D3CP04290A.

[21] F.W. Studier, Protein production by auto-induction in high-density shaking cultures, Protein Expr Purif. 41 (2005) 207–34. doi: 10.1016/j.pep.2005.01.016.

[22] D. Linde, A. Olmedo, A. González-Benjumea, M. Estévez, C. Renau-Mínguez, J. Carro, E. Fernández-Fueyo, A. Gutiérrez, A.T. Martínez, Two new unspecific peroxygenases from heterologous expression of fungal genes in Escherichia coli, Appl Environ Microb. 86 (2020) e02899–19. doi: 10.1128/aem.02899-19.

[23] B. Chance & A.C. Maehly, The assay of catalases and peroxidases, Methods Biochem Anal. 1 (1954) 357–424. doi: 10.1002/9780470110171.ch14.

[24] L.M. Shannon & J.Y. Lew, Peroxidase isozymes from horseradish roots. I. Isolation and physical properties, J Biol Chem. 10 (1966) 2166–2172. doi: 10.1016/s0021-9258(18)96680-9.

[25] M. Girhard, E. Kunigk, S. Tihovsky, V.V. Shumyantseva, V.B. Urlacher, Light-driven biocatalysis with cytochrome P450 peroxygenases, Biotechnol Appl Bioc. 60 (2013) 111–18. doi: 10.1002/bab.1063.

[26] B.O. Burek, S. Bormann, F. Hollmann, J.Z. Bloh, D. Holtmann, Hydrogen peroxide driven biocatalysis, Green Chem. 21 (2019) 3232–49. doi: 10.1039/c9gc00633h.

[27] A. Karich, K. Scheibner, R. Ullrich, M. Hofrichter, Exploring the catalase activity of unspecific peroxygenases and the mechanism of peroxide-dependent heme destruction, J Mol Catal B Enzym. 134 (2016) 238–46. doi: 10.1016/j.molcatb.2016.10.014.

[28] T. Dirks, D. Stoesser, S. Schüttler, F. Hollmann, J. Golda, J.E. Bandow, The atmospheric pressure capillary plasma jet is well-suited to supply H_2_O_2_ for plasma-driven biocatalysis, bioRxiv (2025) 2025.01.07.631711. doi: 10.1101/2025.01.07.631711.

