## Supplementary Information for "Plasma-driven biocatalysis using the cytochrome P450 enzyme CYP152_BSβ_"

**Supplementary Table 1: $CYP152_{BS\beta}$ residual activities after plasma-driven biocatalysis using the capillary plasma jet with 1280 ppm $H_2O$ in the feed gas.**

| | |
|---|---|
| Residual activity after 60 min biocatalysis (Figure 5) [%] | 21.04 ± 8.99 |
| Residual activity after 120 min biocatalysis with frequent buffer exchange (Figure 6) [%] | 8.91 ± 3.03 |

**Supplementary Table 2: Calculations of $CYP152_{BS\beta}$ concentrations and TON using HA403 M beads.**

| **Enzyme loading of beads in immobilization** | |
|---|---|
| amount of beads [mg] | 200 |
| total volume [ml] | 5 |
| $CYP152_{BS\beta}$ concentration [µM] | 10 |
| $CYP152_{BS\beta}$ amount [nmol] | 50 |
| maximum loading of beads [nmol/100 mg beads] | 25 |
| binding efficiency [%] | 96.32 |
| actual loading of beads [nmol/100 mg] | 24.05 |
| **Final concentrations of $CYP152_{BS\beta}$ in reaction** | |
| volume used for reactor [ml] | 1 |
| amount of beads in reactor [mg] | 100 |
| amount $CYP152_{BS\beta}$ in reactor [nmol] | 24.05 |
| reaction volume [ml] | 5 |
| **TON calculations** | |
| product amount [nmol] | 452.59 |
| turnover number | 18.82 |

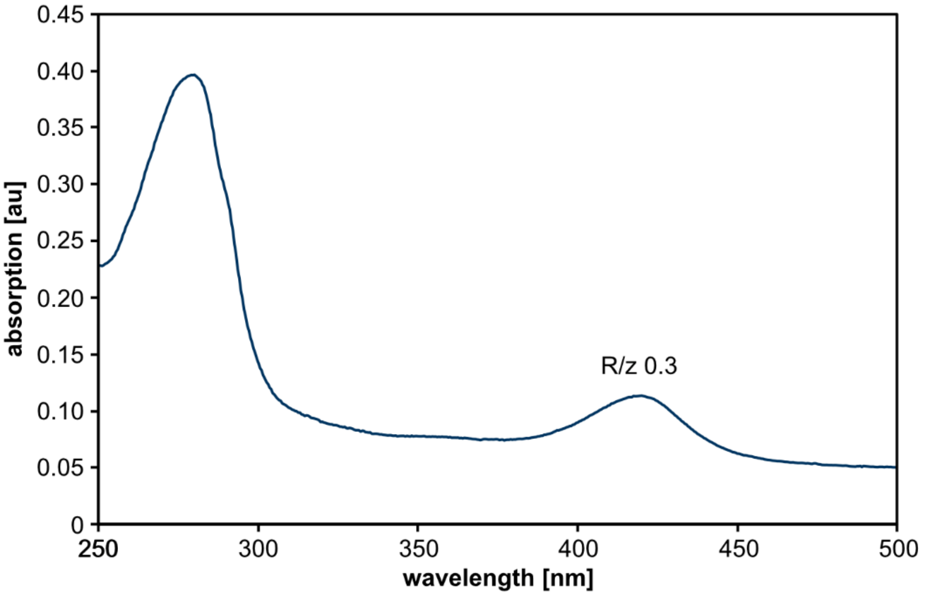


**Supplementary Figure 1: Spectral analysis of purified CYP152$_{BS\beta}$ overproduced in LB medium.** For *cypC* overexpression, cultures were inoculated to an $OD_{600}$ of 0.05 and incubated at 37°C to an $OD_{600}$ of 0.5-0.6, followed by an induction with IPTG (100 µmol $l^{-1}$). After 4 h incubation at 30°C, the cells were harvested and used for protein purification. Absorption spectra were recorded using 12 µmol $l^{-1}$ CYP152$_{BS\beta}$ in potassium phosphate buffer (100 mmol $l^{-1}$, pH 7). Buffer served as blank. R/z value was calculated by relating absorption of the soret peak (at 420 nm) to absorption at 280 nm. Representative data of three independent replicates is displayed.

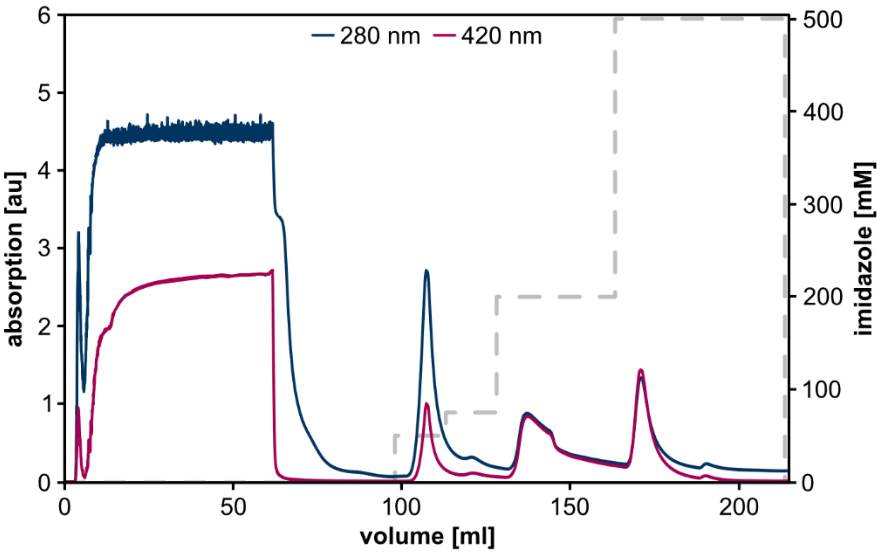


**Supplementary Figure 2: IMAC based purification of CYP152$_{BS\beta}$.** Lysate was loaded onto a HisTrap FF crude 5 ml column (GE Healthcare) and the $His_6$-tagged CYP152$_{BS\beta}$ protein was purified with an ÄKTA pure25 system (GE Healthcare). Proteins were eluted with three stepwise increases in imidazole concentration (50 mmol $l^{-1}$; 75 mmol $l^{-1}$, 200 mmol $l^{-1}$). The HisTrap FF crude 5 ml column was finally subjected to washing with 500 mmol $l^{-1}$ imidazole. Absorption at 280 nm (general proteins) and 420 nm (heme containing protein) is displayed. Representative data of four independent replicates is displayed.

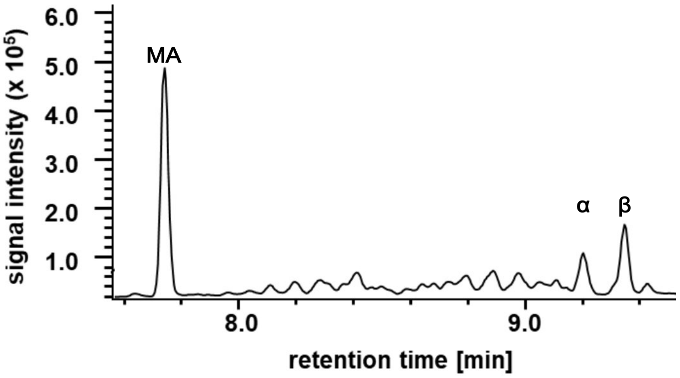


**Supplementary Figure 3: Myristic acid conversion of $CYP152_{BS\beta}$.** The reaction solution consisted of 120 µmol l$^{-1}$ myristic acid, 0.1 µmol l$^{-1}$ $CYP152_{BS\beta}$, and 0.5 mmol l$^{-1}$ $H_2O_2$ in potassium phosphate buffer (100 mmol l$^{-1}$, pH 7). Sample derivatization was performed using BSFTA-TMCS according to Girhard *et al*. [25]. The chromatogram was recorded by gas chromatography analysis (Shimadzu GC2030-Nexis). First peak was identified as myristic acid (MA). Later peaks represented the produced products (α)- and (β)-hydroxy myristic acid. Peak assignment was performed based on recorded mass fragments displayed in Supplementary Figure 3. Representative data of two independent replicates is displayed.

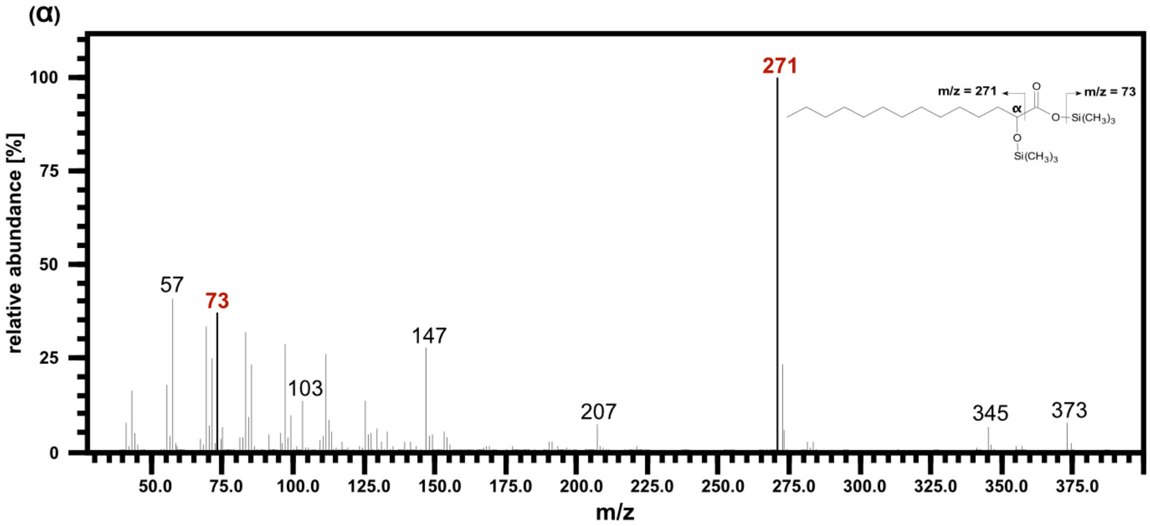


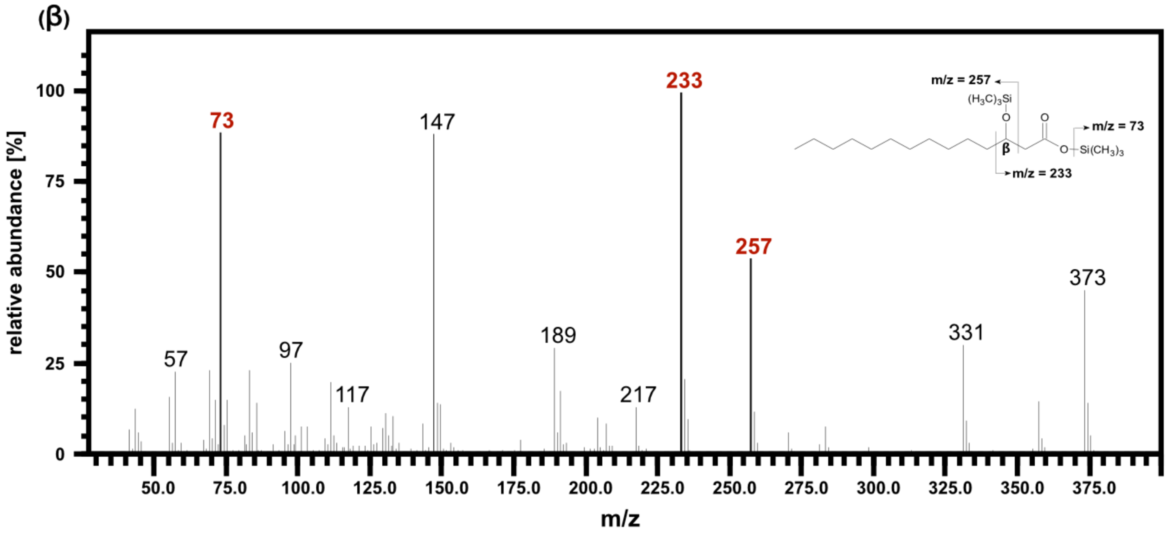

**Supplementary Figure 4: Mass spectra of TMS esters of (α)- and (β)-hydroxy myristic acid after conversion of myristic acid by CYP152$_{BSβ}$.** The reaction solution consisted of 120 µmol $l^{-1}$ myristic acid, 0.1 µmol $l^{-1}$ CYP152$_{BSβ}$, and 0.5 mmol $l^{-1}$ $H_2O_2$ in potassium phosphate buffer (100 mmol $l^{-1}$, pH 7). Sample derivatization was performed using BSFTA-TMCS according to Girhard *et al*. [25]. The chromatogram was recorded by gas chromatography analysis (Shimadzu GC2030-Nexis). Representative data of two independent replicates is displayed.

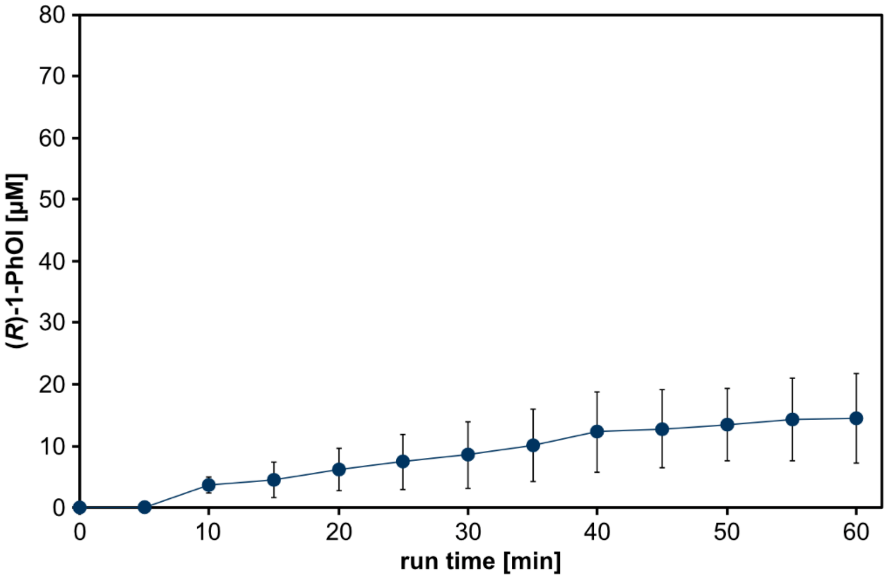


**Supplementary Figure 5: Plasma-driven biocatalysis with capillary plasma jet using CYP152$_{BS\beta}$ and 6400 ppm $H_2O$ in feed gas.** ETBE was hydroxylated using plasma treatment of a reaction solution containing CYP152$_{BS\beta}$ immobilized on ReliZyme HA403 M. The reaction solution contained 5 ml potassium phosphate buffer (100 mmol $l^{-1}$, pH 7) with 50 mmol $l^{-1}$ ETBE and 20 mmol $l^{-1}$ heptanoic acid (as decoy molecule). Plasma treatment was performed with a water concentration of 6400 ppm in the feed gas. Every 5 min aliquots were withdrawn for product analysis by GC measurement. Means and standard deviations reflect three experiments.

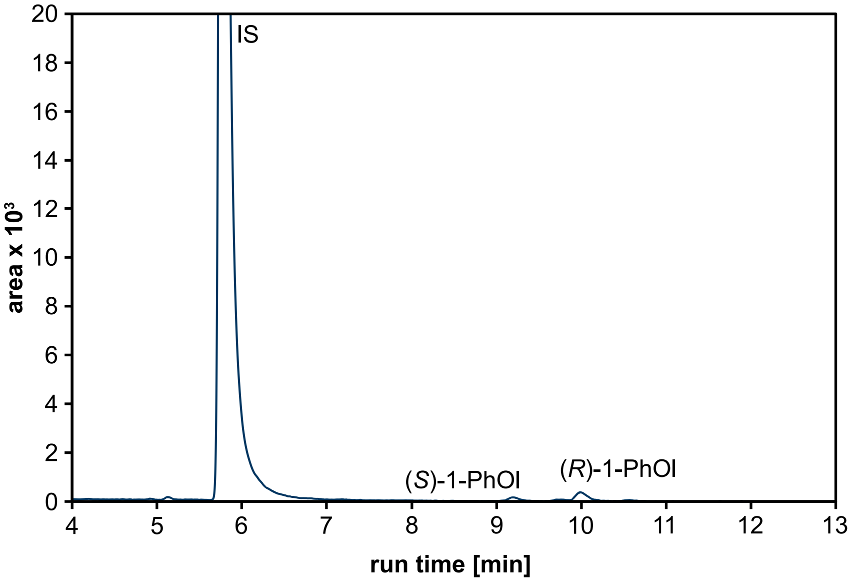


**Supplementary Figure 6: Gas chromatographic analysis of plasma-driven biocatalysis reaction solution using CYP152$_{BS\beta}$.** The substrate ETBE was converted using $H_2O_2$ from plasma treatment of the reaction solution containing CYP152$_{BS\beta}$ immobilized on ReliZyme HA403 M. Reaction solution further contained 5 ml potassium phosphate buffer (100 mmol $l^{-1}$, pH 7) with 50 mmol $l^{-1}$ ETBE and 20 mmol $l^{-1}$ heptanoic acid (as decoy molecule). Plasma treatment was performed with a water concentration of 1280 ppm in the feed gas. IS: internal standard, 1-octanol.

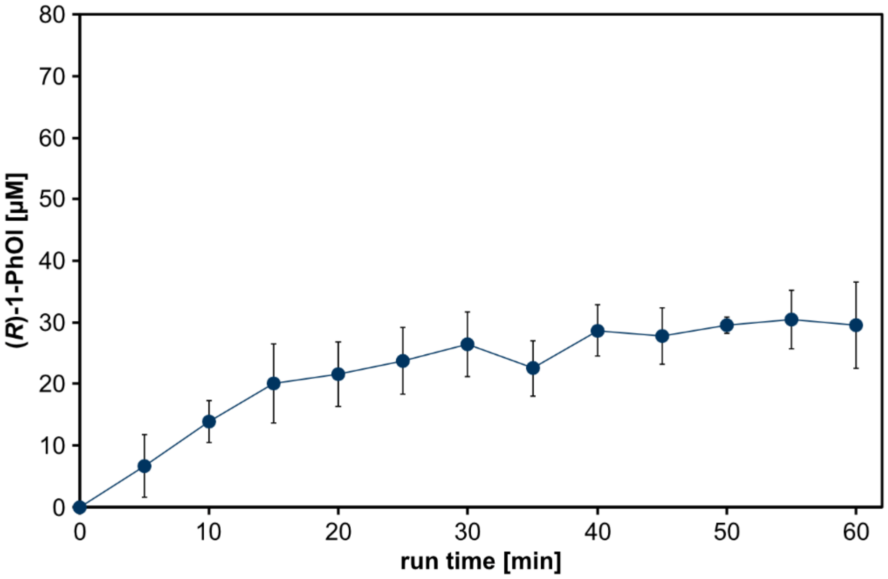


**Supplementary Figure 7: Plasma-driven biocatalysis with capillary plasma jet using CYP152$_{BS\beta}$ and 640 ppm $H_2O$ in feed gas.** ETBE was hydroxylated using plasma treatment of a reaction solution containing CYP152$_{BS\beta}$ immobilized on ReliZyme HA403 M. The reaction solution contained 5 ml potassium phosphate buffer (100 mmol $l^{-1}$, pH 7) with 50 mmol $l^{-1}$ ETBE and 20 mmol $l^{-1}$ heptanoic acid (as decoy molecule). Plasma treatment was performed with a water concentration of 640 ppm in the feed gas. Every 5 min aliquots were withdrawn for product analysis by GC measurement. Means and standard deviations reflect three experiments.
